# From what we perceive to what we remember: Characterizing representational dynamics of visual memorability

**DOI:** 10.1101/049700

**Authors:** Seyed-Mahdi Khaligh-Razavi, Wilma A. Bainbridge, Dimitrios Pantazis, Aude Oliva

## Abstract

Not all visual memories are equal—some endure in our minds, while others quickly disappear. Recent behavioral work shows we can reliably predict which images will be remembered. This image property is called *memorability*. Memorability is intrinsic to an image, robust across observers, and unexplainable by low-level visual features. However, its neural bases and relation with perception and memory remain unknown. Here we characterize the representational dynamics of memorability using magnetoencephalography (MEG). We find memorability is indexed by brain responses starting at 218ms for faces and 371ms for scenes—later than classical early face/scene discrimination perceptual signals, yet earlier than the late memory encoding signal observed at ~700ms. The results show memorability is a high-level image property whose spatio-temporal neural dynamics are different from those of memory encoding. Together, this work brings new insights into the underlying neural processes of the transformation from what we perceive to what we remember.

## Introduction

Recent large-scale behavioral studies show that images carry the attribute of *memorability*, a predictive value of whether a novel image will be later remembered or forgotten (Isola et al., 2011a, 2011b; Bainbridge et al., 2013; Bylinskii et al., 2015). Independent of observers, certain images are consistently remembered and others forgotten (Isola et al., 2011a, 2011b; Bainbridge et al., 2013; Borkin et al., 2013; Isola et al., 2014). Memorability is well-characterized across several stimulus domains, including faces (Bainbridge et al., 2013), scenes (Bylinskii et al., 2015; Isola et al., 2011b), and visualizations (Borkin et al., 2013), and is not explained by simple low-level visual features. Additionally, memorability is shown to be consistent across different local image contexts (Bylinskii et al., 2015) and time lags (Isola et al., 2014). Finally, memorability is image-computable (Isola et al., 2011b; Khosla et al., 2015), and can be modulated by image alterations (Khosla et al., 2013).

Given that memorability is an intrinsic image property, we expect the existence of some neural signatures in the brain discriminating images based on their memorability. These neural signatures are largely unexplored, and may inform our understanding of the greater network of perception and memory. Specifically, is the neural signature of memorability temporally dissociable from those of perception (e.g. Sato et al., 1999; Liu et al., 2002; Rivolta et al., 2012) or memory encoding (e.g. Brewer et al., 1998; Wagner et al., 1998; Rissman et al., 2010; Friedman and Johnson, 2000; Kim, 2011). How might they vary based on stimulus category (e.g. faces, and scenes)? MEG is an ideal technique for addressing these questions, as its high temporal resolution enables us to tease apart processing stages within the order of tens of milliseconds of each other.

Here we reveal a neuromagnetic signature of visual memorability, starting at late stages of perception and before memory encoding. We find this memorability signature for both faces and scenes, with different temporal patterns (a sustained signature over time for faces, and a transient signature for scenes). We further demonstrate that the neural distinction between high memorable and low memorable images exists even within images that are subsequently forgotten–where there is no strong trace of memory. Memorability is thus a distinct neural signature, dissociable from the memory of individuals. This study provides new insights to the long-standing neuroscientific question of how perception selects stimuli for memory encoding and later retrieval.

## Results

### The brain carries information on memorability

In an MEG experiment (Figure S1), human participants performed a perceptual categorization task (indicating male/female for faces; indoor/outdoor for scenes), while viewing 360 novel faces and 360 novel scenes. Unbeknownst to them, the stream of images was composed of high memorable and low memorable images, determined by a large-scale online study (Bainbridge et al., 2013; Isola et al., 2011b) equalized on several image statistics and semantic dimensions (see methods) to eliminate potential confounds with memorability.

To assess whether the MEG signals carry memorability information, we used the representational similarity analysis (RSA) framework (Kriegeskorte et al., 2008a, 2008b; Kriegeskorte and Kievit, 2013; Nili et al., 2014; Cichy et al., 2014; Khaligh-Razavi and Kriegeskorte, 2014) (Figure 1A). This framework enables us to characterize the geometry of a representation in the brain by computing pairwise differences (distances) between patterns of brain activity elicited by a set of stimuli. In contrast, a binary decoding approach (seeking time-points that can classify high memorable versus low memorable images) would not inform on the nature of the representational geometry (Kriegeskorte and Kievit, 2013) of memorability. Thus, we employ a model representational dissimilarity matrix (RDM) that tests specific hypotheses on the shape of the representation. For the case of memorability, behavioral data from our previous studies (Bainbridge et al., 2013; Isola et al., 2011b) showed that high memorable images have significantly different mean memorability scores than low memorable images, yet their individual standard deviations are similarly small. This suggests a neural representational space where stimuli are tightly clustered based on their level of memorability. To search for time-points in the MEG signal sensitive to this representational space, we created a reference model RDM, in which images of high memorability were assumed similar to each other, and distinct from low memorable images; and vice versa (Figure 1A). We refer to this reference RDM as the memorability model RDM. We also explored two other possible representational geometries for memorability (see Figure S2).

**Figure 1.**
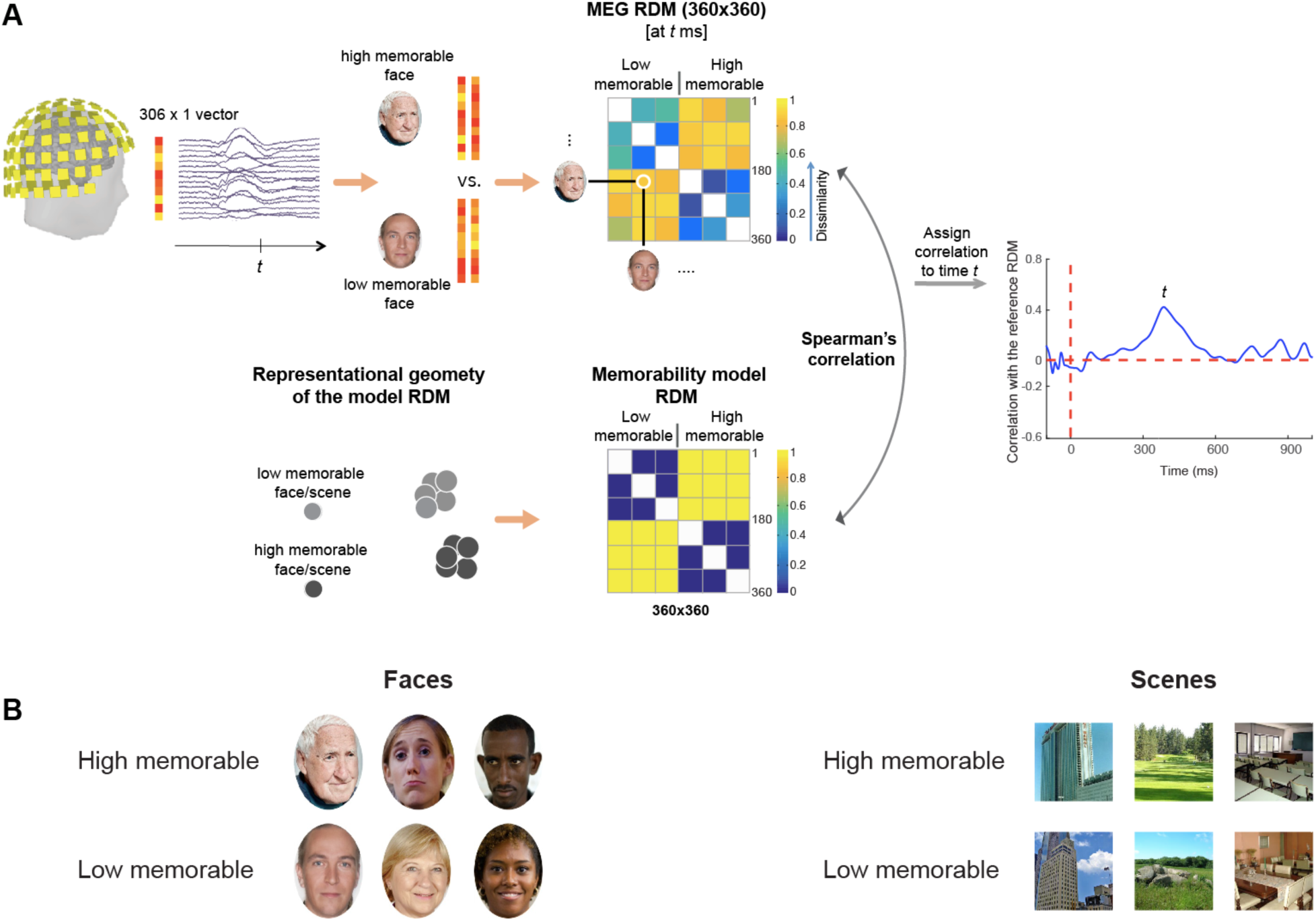
Representational similarity analysis of MEG data. A) Estimation of face and scene memorability neural representations. For each time point, we constructed a 360x360 MEG RDM by computing the pairwise dissimilarities (one minus Pearson correlation) between the MEG patterns of all face images. Each MEG RDM was then compared (Spearman’s rank correlation) against a memorability model RDM with elements set to 1 between high versus low memorable images, and 0 within high or within low memorable images. Correlations were then normalized relative to the noise-ceiling (see methods), resulting in a face memorability time series (Figure 2A). Same analysis was repeated for scene memorability (Figure 2B). B) Example face and scene images. We presented a total of 360 faces and 360 scenes. In each category half of the images had high memorability and the other half had low memorability. Face images were controlled for gender, race, age, emotion, attractiveness, friendliness, confidence, false alarm rate, spatial frequency and color. Scene images were controlled for indoor/outdoor, natural/manmade, no people or animals, false alarm rate, spatial frequency, color, number of objects, and size of objects. The stimuli shown in this figure are for illustrative purposes, and available to the public domain (Creative Commons License).

**Figure 2.**
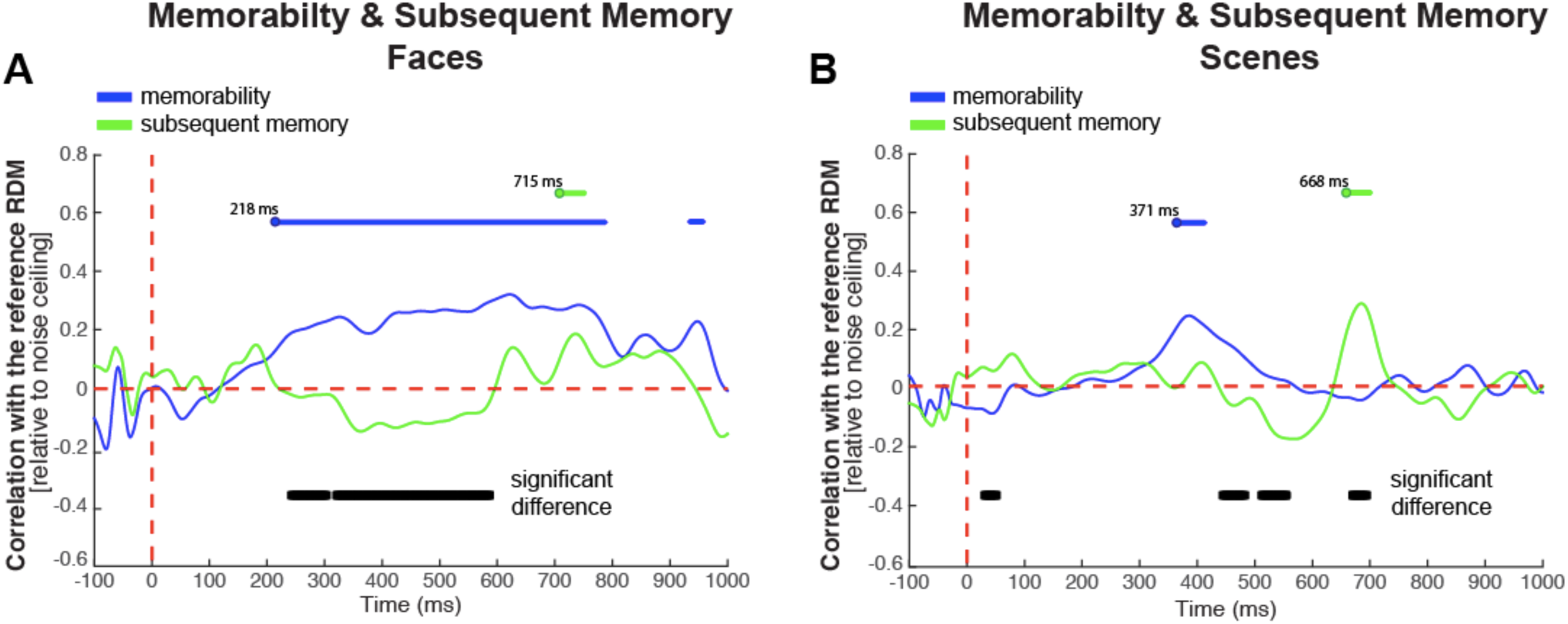
Time courses of memorability and subsequent memory. Time courses of memorability representations for faces (A) and scenes (B) were constructed by correlating their respective MEG RDMs at each time-point with the reference memorability RDM, as illustrated in Figure 1. Face memorability had earlier onset (218 ms) than scene memorability (371 ms) and was more sustained over time. Following the MEG perceptual categorization experiment, we administered an unanticipated memory recall behavioral task, results of which were used to construct subject-specific (face or scene) memory RDMs, with elements set to 1 between remembered versus forgotten images, and 0 for within remembered or forgotten images. Time courses of subsequent memory representations were then constructed similar to the memorability representations, but comparing the MEG data against subject-specific memory RDMs rather than the memorability model RDM. Memorability (blue line) and subsequent memory (green line) representations have different temporal dynamics for both faces (A) and scenes (B). Blue and green horizontal lines above the curves indicate significant memorability and subsequent memory time points, respectively (10,000 permutations of condition labels, false discovery rate corrected at 0.05). Horizontal black lines below the memory/memorability curves indicate time points with significant difference between the two curves (two-sided signed-rank test, false discovery rate corrected at 0.01). Note, an observed early difference (at 31 ms) between memory encoding and memorability in scenes is a spurious result since there is no significant memory or memorability signal so early in time.

This memorability model RDM was then compared to RDMs constructed from the MEG data to examine if at any time-points brain signals carry information about image memorability. From the MEG neural signals, we created two 360x360 RDMs at each time-point (from 100ms before stimulus onset to 1000ms after stimulus onset), one for face stimuli and another for scenes. The MEG RDMs represent the pairwise dissimilarity between the MEG signals elicited by either the 360 face stimuli or the 360 scene stimuli (see Figure 1B for sample stimuli). To assess the ability of MEG signals in discriminating high memorable versus low memorable images, we then correlated the MEG RDMs with the memorability model RDM across time. We found that for both faces and scenes, the memorability model RDM explained a significant non-noise variance of the MEG signals across time (Figure 2A, B). This demonstrates that the brain carries information about memorability, and that we can determine from the MEG signals whether the observed stimulus has high memorability or low memorability.

While we find a clear memorability signature for both faces and scenes, their particular representational patterns differed. Faces showed a persistent memorability signature, with an early 218 ms signal onset and a sustained representation (up to ~550ms). In contrast, scenes showed a transient memorability signature, with a late 371 ms onset and a much shorter signal duration (~100ms). Overall, the results suggest that our brain carries a distinct signal of memorability for different stimulus categories.

### Memorability is a late perceptual signal, and temporally dissociable from the subsequent memory signal

Previous large-scale memorability studies (Bylinskii et al., 2015; Khosla et al., 2015), have shown that memorability is a multi-facet high-level image property. That is, we can think of memorability as a combination of several other high-level image properties (e.g. popularity, visual distinctiveness, emotion, abnormality), each of which partially contributes in explaining some variance of image memorability.

To compare the neural signatures of this multi-facet image property (memorability) against the temporal dynamics of memory encoding (Kim, 2011; Friedman and Johnson, 2000), following the MEG recording, the participants were asked to complete an unanticipated memory test evaluating which images were eventually remembered or forgotten (see Methods). For each participant, we then summarized the results of their subsequent memory test in a model RDM, designed to have pairwise dissimilarity values between forgotten and remembered images set to 1, and within forgotten or within remembered images set to 0. These matrices are called subsequent memory model RDMs. We then correlated the MEG RDMs of each participant with their subsequent memory RDMs at each time-point (Figure 2, green curve), which gave us the subsequent memory signal (also called memory encoding through out the paper). The subsequent memory signals, as opposed to the memorability signals, showed a significant effect starting very late in time (715ms for faces and 668ms for scenes) [random permutation test, FDR-corrected across time].

Critically, we examined whether the signatures of memory encoding and memorability differed across time. Indeed, for faces we found a significantly stronger neural signature of memorability compared to subsequent memory between 243 to 589 ms, indicating that brain signals capture memorability better than subsequent memory (Figure 2A) [two-sided sign-rank test, FDR-corrected at 0.01]. For scenes, however, we found a pattern of significant differences with memorability being higher than subsequent memory and vice versa at different time points (Figure 2B).

Even though the temporal signatures of memorability and subsequent memory were different, memorability and subsequent memory RDMs were also significantly correlated (Figure S3). Indeed, by the definition of memorability, we expect high memorable images to be remembered more often than low memorable ones. However, one effect does not fully explain the other. Memorability is an image property independent of an individual observer, while memory is an observer-specific process. As there are time-points where memorability effect is significantly more correlated with brain responses than subsequent memory and vice versa (Figure 2B), memorability is not just a proxy for memory encoding, nor is memory encoding just a noisier version of memorability.

In sum, for both faces and scenes we observed that neural signatures of memorability significantly differed from those of subsequent memory. This suggests that memorability information carried by the brain is different from subsequent memory information, therefore emphasizing that they have different underlying neural processes. However, they are also related processes and work together as part of the larger network of perception and memory. Potentially, the function that memorability serves might be guiding the storage of information into memory.

### Spatial localization of memorability

Given the reasonably good spatial resolution of MEG, we also examined a sensor-wise visualization of memorability effect. While memorability may show largely separate temporal patterns from early perception and memory encoding, do these differences also emerge in terms of their spatial patterns? With this, we can approach a spatio-temporally resolved understanding of memorability. For subsequent memory, we would predict activations in regions implicated in previous subsequent memory work, specifically temporal, parietal, and frontal regions (Fernández and Tendolkar, 2001; Eichenbaum et al., 2007; Kim, 2011). For memorability, we may predict activations in occipital and temporal perceptual regions, as well as possibly in a subset of regions implicated by memory.

For scenes, where we had transient temporally non-overlapping patterns of memorability and memory, we provided a sensor-wise visualization at the peak latency of the observed experimental effects (Figure 3). Posterior and temporal sensors were involved in carrying scene memorability information, while the subsequent memory effect was more distributed and further extended towards the frontal and central sensors, and was more left-lateralized.

**Figure 3.**
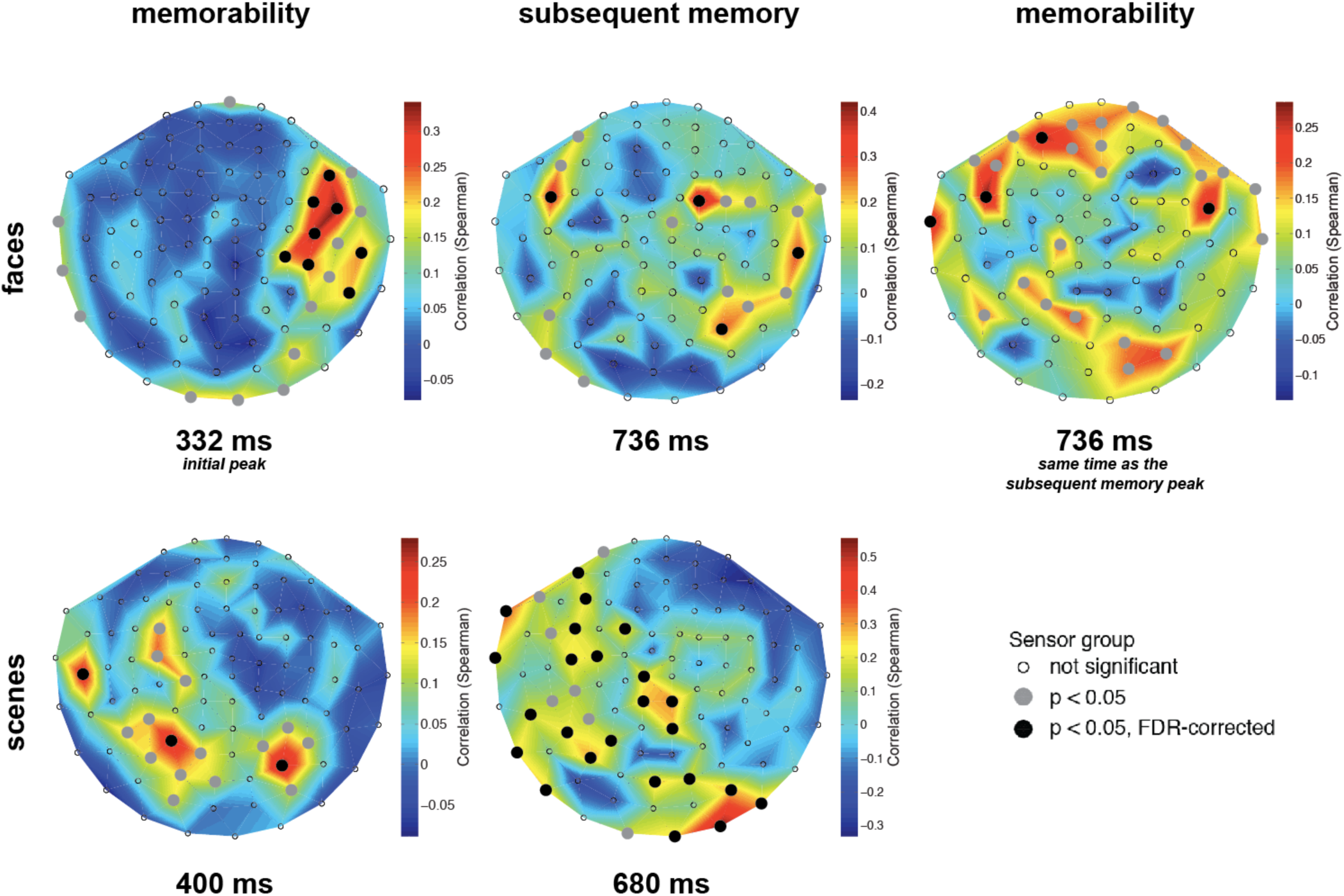
Sensor-wise RDM correlation maps at the peak latency of observed experimental effects. To investigate which sensors contributed to observed experimental effects, we conducted an equivalent RSA analysis for each MEG channel (306) separately at the peak latency of the effects in question. Since MEG sensors were spatially arranged in 102 triplets (Elekta Triux system; 2 gradiometers and 1 magnetometer in each location), we computed dissimilarity matrices for triplets and compared them with the reference model RDM (for memorability or subsequent memory), yielding 102 data points. Sensor-wise visualization of face memorability effect is shown at two time-points: an initial peak at 332 ms, and a later time-point at 736 ms when the memorability effect overlaps with the subsequent memory peak. Sensor-wise visualization of scene memorability is shown at its peak (400 ms), compared with the peak of scene subsequent memory effect at 680 ms. Significant sensors are shown with black dots (random permutation test, 10,000 permutations; FDR-corrected across sensors). The overall sensor-wise patterns between memorability and subsequent memory suggest that each effect is driven by different sources in the brain.

For faces, where we had a sustained memorability effect, Figure 3 shows sensor-wise visualizations at two time-points: the initial peak at 332 ms, and a later time-point at 736 ms, which is the same as the observed peak for subsequent memory effect. For the initial peak at 332 ms, a weak memorability effect was present in posterior sensors and a significant effect in the right temporal sensors. The later face memorability at 736 ms, which is overlapping with the peak of subsequent memory effect, was more fronto-temporal, yet different from the sensor-wise pattern observed for memory encoding at the same time.

Further to this, to study the spatial sources of the sustained face memorability effect over time, we visualized the sensor-wise memorability effect over the time-window in which face memorability was observed [218 ms to 796 ms] (Figure 4, Supplementary Movie 1). This enabled us to see that the face memorability information travels through different brain areas over time, and the observed sustained activity is not the result of one brain area constantly responsive to face memorability. The neural signature of face memorability starts from right posterior sensors (at 218 ms) and gradually moves to the right temporal sensors (280 to 500 ms). It then reaches the frontal sensors at 550 ms to 600 ms; after which the signal becomes more distributed and travels to the left temporal sensors and is weakly present in some posterior sensors too.

**Figure 4.**
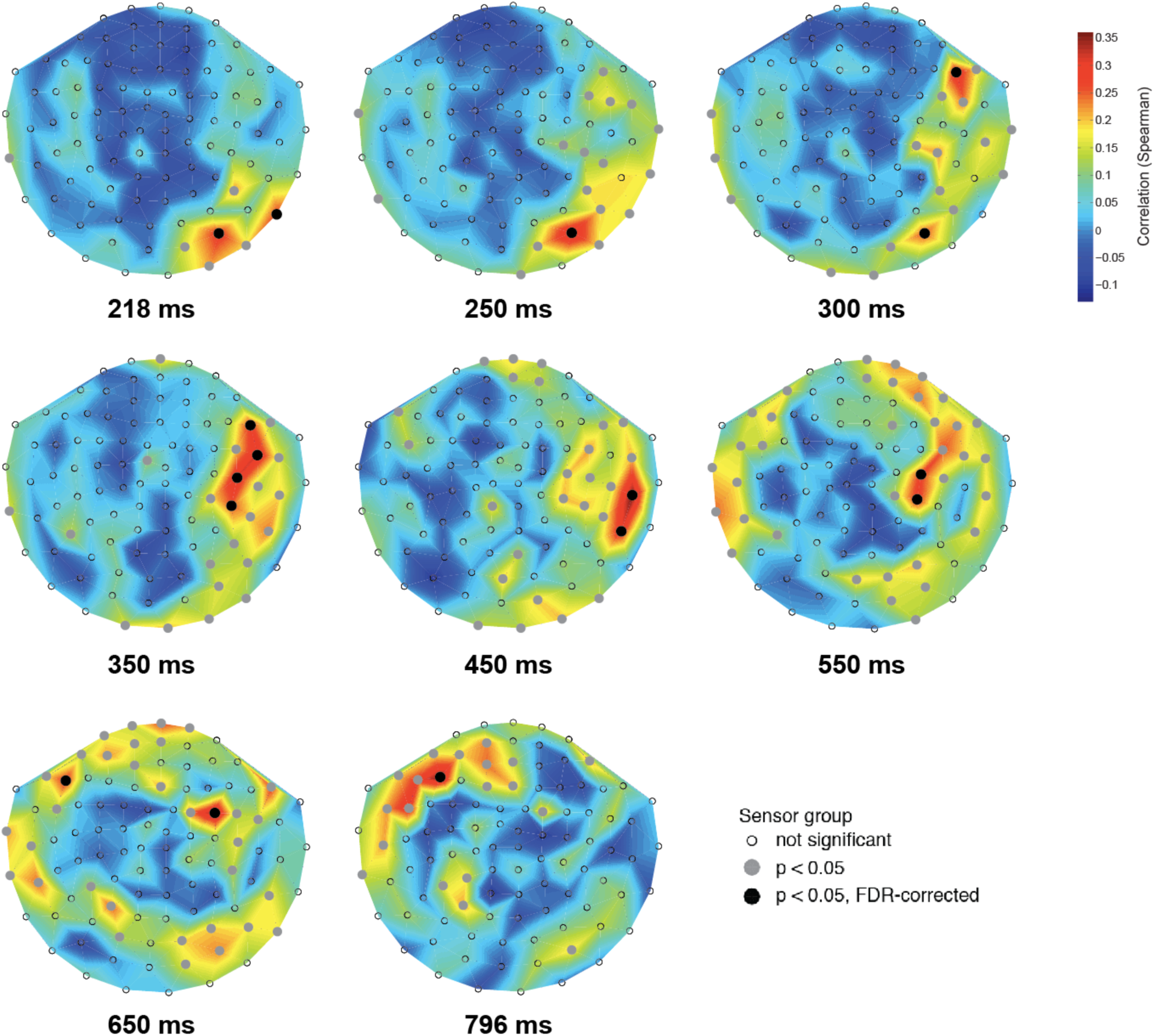
Sensor-wise visualization of face memorability over time. Faces elicited a sustained memorability effect from 218 ms to 796 ms, with spatial patterns varying across time indicating the contribution of a cascade of cortical regions. Sensor-wise maps were obtained as in Figure 3. FDR correction for multiple comparisons was performed for time and sensors. See Supplementary Movie 1 for all time-points.

Overall, we find that face memorability is more right lateralized compared with scene memorability; and face memory seems spatially more distributed than the scene memory, which is more left lateralized.

The sensor-wise analysis revealed that for scenes and earlier face memorability (before 500 ms), several visual regions (posterior and temporal sensors) were involved. This is consistent with memorability being a high-level perceptual signal. For later face memorability signals, some central and frontal sensors were also engaged–which may be more related to memory regions (Fernández and Tendolkar, 2001). This is also expected given that memorability is not totally unrelated to memory (as mentioned earlier), and that there is a continuum (Bussey and Saksida, 2007) between perception and memory. In this continuum, if the role of memorability is to link perception and memory, we would expect to see temporal cortex–where visual perception meets memory (Miyashita, 1993; Murray et al., 2007)— to be involved. Consistent with this assumption, we observed that temporal sensors were substantially involved in both face and scene memorability.

### Memorability of forgotten images

If memorability is a true phenomenon, separate from subsequent memory signals (Brewer et al., 1998; Friedman and Johnson, 2000; Wagner et al., 1998), we should be able to see a trace of memorability within forgotten images, where there is no detectable memory effect.

To this end, for each individual, we re-analyzed their MEG data for the memorability effect (similar to the procedure explained in Figure 1) including only images that were subsequently forgotten by a given participant. We then averaged the results across all participants.

The results (Figure 5) revealed that the memorability effect is present even within forgotten images, with a particularly strong effect for faces (Figure 5A). Note that the weaker effect for scenes may be due to the fact that overall fewer scenes were forgotten compared to faces—so we had less data.

**Figure 5.**
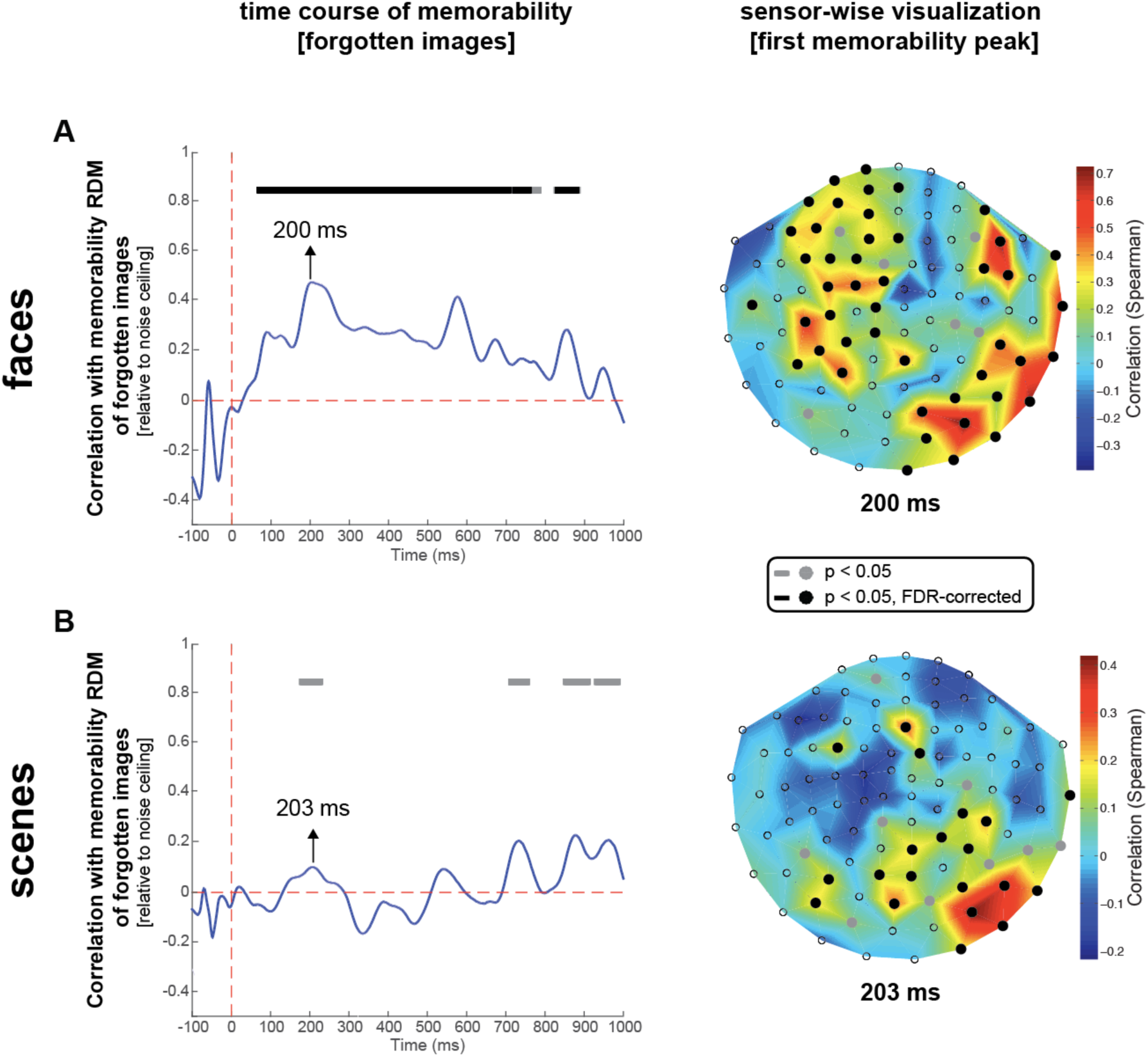
Memorability signature exists even within subsequently forgotten images. For each participant, we repeated analysis in Figure 2 but constrained to the images that were forgotten in the subsequent memory experiment. We also visualized the sensor-wise correlation maps of the memorability signals in the first memorability peak. This indicates the spatial sources indexing memorability of forgotten images. Gray dots/lines indicate significant memorability effect (p < 0.05) before correction for multiple comparisons (random permutation test, 10,000 permutations). Black dots/lines indicate significant effect after FDR correction (corrected across time for the plots on the left, and corrected across sensors for the sensor-wise visualizations on the right).

The memorability of forgotten images starts earlier in time, and based on sensor-wise visualizations (Figure 5), it seems that here more ventral-stream regions are involved (more posterior and temporal sensors) than the previous analysis, where all images were included. This is expected given that the measured memorability effect here is orthogonalized to memory (more of a perceptual signal not contaminated by memory). For face memorability it seems that in addition to the engagement of more ventral-visual-stream regions, some frontal regions are also involved. This is consistent with the existence of an extensive network of face processing regions (Haxby et al., 2000; Ishai et al., 2005), including regions in the frontal lobe (Tsao et al., 2008a, 2008b; Baker, 2008).

The fact that a memorability signature exists even for forgotten images, identifies memorability as a high-level perceptual signal that can be characterized independent of memory.

## Discussion

Humans are very good at remembering images (Brady et al., 2008; Konkle et al., 2010), but not all images are equally memorable. Some images are better remembered (high memorable) and some are quickly forgotten (low memorable).

We demonstrated that memorability is a high-level perceptual property of an image, which is content-dependent. That is the representational dynamics of memorability change with the stimulus type (face vs. scenes). Yet we showed that for both faces and scenes, the onset of memorability is after early stages of face/scene perception (Liu et al., 2002; Rivolta et al., 2012; Sato et al., 1999), and before memory encoding. Furthermore, memorability and memory had different sensor-wise MEG patterns, suggesting they have different spatial sources in the brain. Importantly, the neural signature of memorability existed even within subsequently forgotten images. Taken together, these results provide evidence for a novel neuromagnetic signal of higher-level perception, dissociable from subsequent memory signal, that can be thought of as an intermediate stage of processing that links perception to memory.

### Memorability links perception to memory encoding

The memorability effects cannot be explained by low-level visual differences or memory encoding. Face perception is known to start early after stimulus onset at ~100 ms (Bentin et al., 1996; Linkenkaer-Hansen et al., 1998; Halgren et al., 2000; Liu et al., 2002; Cauchoix et al., 2014), and scene perception is known to start at about ~200 ms (Sato et al., 1999; Vanrullen and Thorpe, 2001; Rousselet et al., 2005; Rivolta et al., 2012). Here we observed that for both faces and scenes the model of memorability outperforms that of memory encoding in correlations with neural activity only later in time (starting at 243 ms after stimulus onset for faces, and at 445 ms for scenes), suggesting that the observed differences in the time-courses of memory and memorability are not due to low-level visual differences. Indeed, the stimuli used in this study were highly controlled for low-level visual differences (see Methods), so we expected no early neural differences due to low-level image properties to emerge related to memorability. Furthermore, we observed that later in time the subsequent memory signal gets significantly stronger than the memorability signal for scenes, and we found different spatial loci for patterns of memorability and subsequent memory. These results indicate that the observed subsequent memory effects are not just noisier measures of memorability.

Given that the observed differences between memorability and memory encoding time-courses are neither due to low-level visual differences, nor due to a higher noise-level in the memory signal, but a genuine effect observed both temporally and spatially, it would be interesting to know how the two signals are related to each other and to perception. We found that the memorability neural signal for both faces and scenes started well after initial face or scene perception (~[100 to 200 ms] see (Sato et al., 1999; Liu et al., 2002; Rivolta et al., 2012)) and before the start of the memory encoding signal observed here, and reported by previous works (400-800 ms (Paller et al., 1988; Friedman and Johnson, 2000)). The temporal patterns we observed here, as well as the prevalence of memorability effect in temporal sensors (where perception and memory meet (Miyashita, 1993; Bussey and Saksida, 2007; Murray et al., 2007)), suggest that memorability is an intermediate processing stage, linking perception to memory.

### Sustained face memorability vs. transient scene memorability

Faces showed an early, persistent memorability signature whereas scenes showed a later, transient memorability signature. Why might these content-based differences emerge in the memorability signal? Face stimuli are important for social interaction, they attract more attention (Vuilleumier, 2000; Devue et al., 2009; Gliga et al., 2009), and there exists an extensive network of areas involved in face processing (Kanwisher et al., 1997; Haxby et al., 2000; Rossion et al., 2003; Ishai et al., 2005; Baker, 2008), which may help maintain a persistent face signal over time. From our sensor-wise visualization of face memorability (Figures 4), we observed that several brain regions are engaged in carrying face memorability information over time, leading to a memorability signal persisting longer in the brain–being carried from one region to another. In the case of scene stimuli, fewer sensors were involved in carrying scene memorability information across the brain (see Figure 3), resulting in a transient memorability effect for scenes. This may occur because scenes are on average more memorable than faces (Isola et al., 2011b; Bainbridge et al., 2013), so differences between low and high memorable scenes may be smaller and trigger less of a pattern difference. Regardless, these different characteristics of face and scene memorability show that while memorability effects exist across different content domains, perception, memorability, and memory encoding may interact differently depending on content type. Further investigation will be needed to see what aspects of memorability are content-specific, and whether other differences emerge in other domains such as objects or words.

## Experimental Procedures

### Stimuli

The stimulus set comprised 720 images, equally divided into four conditions: 1) low memorable faces, 2) high memorable faces, 3) low memorable scenes, and 4) high memorable scenes. Memorability was determined for all images based on hit rates (HRs) from an online crowd-sourcing study on Amazon Mechanical Turk, as described in detail in (Isola et al., 2011b).

The 360 faces images were taken from the 10k US Adult Faces Database, which includes memorability scores, demographic information, and attribute labels for 2,222 faces (Bainbridge et al., 2013). High memorable faces were taken from the top 25% of the HR distribution (M=0.72, SD= 0.07), and low memorable faces were taken from the bottom 25% (M=0.32, SD=0.05), with both sets significantly different in HR (*t*_(358)_ = 62.01, *p* < 10^−6^), but not significantly different in false alarm rate (FAR) (*t*_(358)_ = 1.30, *p* = 0.196). To control for low-level visual and other confounds, face images were balanced across gender, race, age, emotion, attractiveness, friendliness, confidence, false alarm rate, spatial frequency and color. Thus, each condition was 50% female and 50% male. Race roughly followed the distribution of the US population (U.S. Census Bureau, 1990) for both conditions (85% white, 8% black, 5% Hispanic, 5% Asian). Facial expression was balanced between conditions, with both having a distribution of 68% happy, 31% neutral, and 1% sad faces. There was no significant difference in perceived age (*t*_(356)_ = 0.88, *p* = 0.380) between conditions. Additionally, there was no difference in rated coldness, confidence, introversion, kindness, attractiveness, friendliness, emotional magnitude, or happiness (all p > 0.1) between conditions. Lastly, no difference in spatial frequency information was found between conditions (Torralba and Oliva, 2003).

The 360 scene images were taken from the SUN Database (Xiao et al., 2010), with memorability and attribute information from (Isola et al., 2011b). As with faces, high memorable scenes were from the top 25% of the HR distribution (M=0.98, SD = 0.02) and low memorable scenes were from the bottom 25% (M=0.69, SD=0.08). HR was significantly different between these two groups (*t*_(358)_ = 47.11, *p* << 10^−6^), while FAR was not (*t*_(358)_ = 1.96, *p* = 0.051). The two conditions did not have significantly different spatial frequency information (Torralba and Oliva, 2003), and also had no differences in mean or variance in color in both RGB and Lab space (p > 0.05 for all). Each condition contained 50% indoor and 50% outdoor scenes, with the outdoor scenes being 42% natural and 58% manmade. Scenes were chosen so they did not contain any people, animals, central objects, or text. No difference was found between conditions in aesthetics ratings, number of objects, average object size, or maximum object size (all p > 0.1).

Foil faces and scenes for the post-MEG memory task were selected to have similar parameters to the above target stimuli shown during the experiment. They were selected based on the same criteria in terms of stimulus properties sub-categories, except they were selected from the middle 50% of HR. Target and foil faces had no differences in gender, race distribution, false alarm rate, spatial frequency information, or higher-level attributes (e.g. attractiveness). Target and foil scenes had no differences in indoor/outdoor, manmade/natural, spatial frequency information, or color statistics.

### Participants and MEG experimental design

Fourteen right-handed, healthy volunteers with normal or corrected-to-normal vision (7 female, age mean ± s.d. = 24.28 ± 5.5 years) participated in the experiment. The study was conducted according to the Declaration of Helsinki and approved by the local ethics committee (Institutional Review Board of the Massachusetts Institute of Technology). Informed consent was obtained from all participants.

During MEG recordings participants completed an orthogonal image categorization task. Participants viewed a sequence of images (6° visual angle; 0.5 sec duration) while fixating on a cross presented at the center of the screen. Every 2-4 trials (randomly determined), a question mark appeared on screen prompting participants to perform a two alternative forced-choice task between male/female if the last stimulus was a face, or indoor/outdoor if the last stimulus was a scene. This task was designed to ensure participants were attentive to the images without explicitly encoding them into memory. Participants only responded in selected trials to prevent contamination of MEG signals with motor activity. The experiment was divided into 16 runs, with each run containing 45 randomly selected images (each of them shown twice per run—not in succession), resulting in a total experiment time of about 50 min.

### MEG acquisition

We acquired continuous MEG signals from 306 channels (204 planar gradiometers, 102 magnetometers, Elekta Neuromag TRIUX, Elekta, Stockholm) at a sampling rate of 1 kHz, filtered between 0.03 and 330 Hz. Raw data were preprocessed using spatiotemporal filters (Taulu and Simola, 2006; Taulu et al., 2004) (maxfilter software, Elekta, Stockholm) and then analyzed using Brainstorm (Tadel et al., 2011). MEG trials were extracted with 100 ms baseline and 1000 ms post-stimulus, the baseline mean of each channel was removed, and the data were temporally smoothed by a 15Hz low-pass filtering.

### Subsequent memory test

After the recording session, participants were asked to complete an unanticipated memory test, to obtain which images were ultimately remembered or forgotten. Participants viewed a sequence of randomly-ordered face and scene images, half from the MEG study, and half matched foil images. Images were presented for 1s, but the response period was self-paced, and participants had to press a button indicating whether each image was already seen (in the previous MEG experiment) or unseen (new) to proceed to the next trial. This test was divided into four blocks and lasted approximately 40min. As expected, high memorable stimuli were better remembered than low memorable stimuli (Figure S4). Note that the relatively low memory recognition performances are expected, given that half of the images were purposely designed to be forgettable. Furthermore, subjects were not expecting to be tested on their memory (implicit memory task); this further makes the task difficult. Nevertheless, the reported behavioral performances here are comparable to that of Brewer et al., (1998)–the first neuroimaging study of subsequent memories.

### Representational similarity analysis (RSA)

RSA enables us to relate representations obtained from different modalities (e.g. computational models, MEG and fMRI patterns) by comparing the dissimilarity patterns of the representations (Kriegeskorte et al., 2008a, 2008b; Kriegeskorte and Kievit, 2013; Nili et al., 2014; Cichy et al., 2014; Khaligh-Razavi and Kriegeskorte, 2014). In this framework, a representational dissimilarity matrix (RDM) is a square symmetric matrix in which the diagonal entries reflect comparisons between identical stimuli and are set to 0, by definition. Each off-diagonal value indicates the dissimilarity between the activity patterns associated with two different stimuli. In this study, the MEG response patterns at time point t evoked by the different images were compared to each other using representational dissimilarity matrices (RDMs). The measure for dissimilarity was correlation distance (1- Pearson linear correlation) between the MEG signals (from 306 channels).

Given that each stimulus was presented twice, for each of the two categories (faces, scenes), we constructed two RDMs at each time-point: 1) pairwise dissimilarities between the first presentation of the stimuli, 2) pairwise dissimilarities between the second presentation of the stimuli. We found no significant differences between the correlations of the two RDMs (1^st^ and 2^nd^ presentation) with our reference RDMs (memorability model RDM, or subsequent memory RDM) across time (signed-rank test, FDR corrected at 0.01). Therefore, to reduce the noise, for each stimulus, we averaged the MEG patterns of the two presentations and then built the RDMs by calculating the pairwise dissimilarities between the averaged responses. The reported RDM correlations (MEG-memorability, and MEG-memory) are based on the averaged results.

#### Subsequent memory RDM

this is a model RDM based on a post-MEG memory recognition test. The pairwise dissimilarity values within all the remembered images were set to 0, the pairwise dissimilarity values within all the forgotten images were set to 0; and the pairwise dissimilarity values between forgotten and remembered images were set to 1. We construct two subsequent memory RDMs for each subject, one for faces, and one for scenes.

#### Sensor-wise visualizations

For selected time-points, to investigate which sensors contributed to observed experimental effects, we conducted an equivalent RSA analysis for each MEG channel separately. There were 306 MEG sensors that were spatially arranged in 102 triplets (Elekta Triux system; 2 gradiometers and 1 magnetometer in each location). We thus computed dissimilarity matrices for each group of sensors (102 groups of 3 sensors) and correlated them with their respective reference model RDM (i.e. memorability RDM, or subsequent memory RDM), yielding 102 data points that were then visualized.

### Significance testing

We used non-parametric statistical tests, which do not make assumptions about the distribution of the data, for random-effects analysis. For inferential analysis that involved testing whether two patterns are significantly different over time (e.g. comparing temporal patterns of memory vs. memorability), we used two-sided Wilcoxon signed-rank test, which is a non-parametric statistical test (Gibbons and Chakraborti, 2011; Hollander and Wolfe, 1999). For one-sided non-paired tests (i.e. significant RDM correlations), we used random permutation test of conditions. Random permutation tests were based on 10,000 randomizations of the condition labels. When testing memorability, the condition labels were memorable/forgettable; when testing memory, the labels were remembered/forgotten. To correct for multiple comparisons, where applicable, we used the false discovery rate (FDR) (Benjamini and Hochberg, 1995; Groppe et al., 2011). FDR-controlling procedures control the expected proportion of incorrectly rejected null hypotheses (“false discoveries”) among all rejected null hypotheses.

#### Noise ceiling

To account for limits on the amount of dissimilarity variance a model RDM could explain due to noise in brain recordings, we normalized by the noise-ceiling. For each time point, we defined the noise-ceiling as the average pairwise Spearman’s rank correlation between the fourteen RDMs (fourteen subjects). This was done separately for faces and scenes. We then normalized the MEG-to-model RDM correlations relative to the noise-ceiling.

## Acknowledgments

We would like to thank Santani Teng Radoslaw Martin Cichy, Zoya Bylinskii and Caitlin Mullin for helpful comments and discussions. This work was co-funded by the *McGovern Institute Neurotechnology Program* and NSF award 1532591 in *Neural and Cognitive Systems* (to A.O and D.P.). A.O was also partly supported by the Guggenheim foundation. The study was conducted at the Athinoula A. Martinos Imaging Center at the McGovern Institute for Brain Research, Massachusetts Institute of Technology.

---

**Figure S1.**
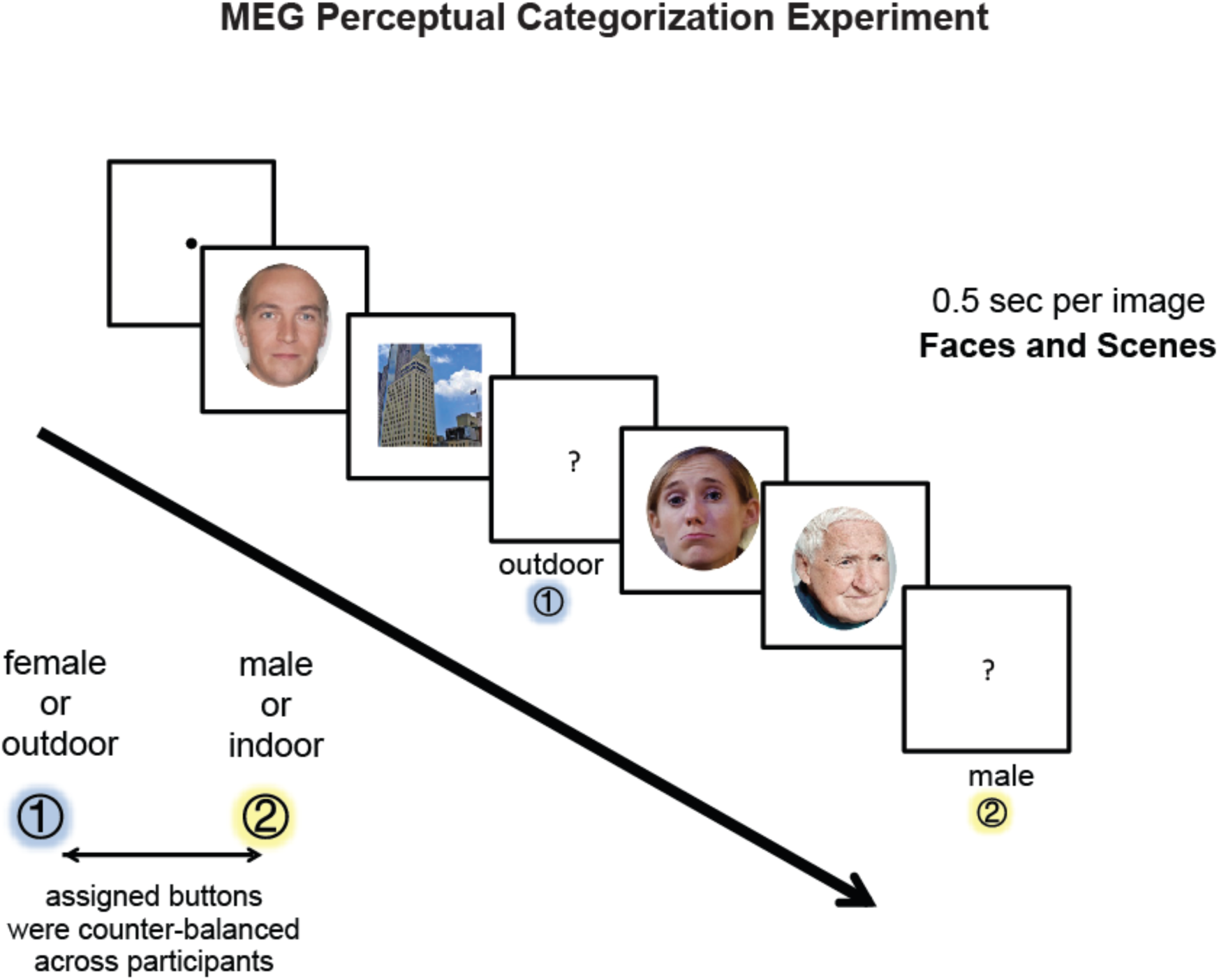
(related to ‘Experimental Procedures’). **MEG Experiment** there were 360 faces and 360 scenes. In each category half of the images had high memorability and the other half had low memorability. Face images were controlled for gender, race, age, emotion, attractiveness, friendliness, confidence, false alarm rate, spatial frequency and color. Scene images were controlled for indoor/outdoor, natural/manmade, no people or animals, false alarm rate, spatial frequency, color, number of objects, and size of objects. Images were presented in random order and were on for 0.5 sec. Every 2 – 4 trials, a question mark was presented prompting a button press response so participants can indicate the category of the last image (male/female, indoor/outdoor). The stimuli shown here are for illustrative purposes, and available to public the domain (Creative Commons license). Link to the stimuli: faces (http://www.wilmabainbridge.com/facememorability2.html) and scenes (http://web.mit.edu/phillipi/Public/MemorabilityPAMI/index.html).

**Figure S2.**
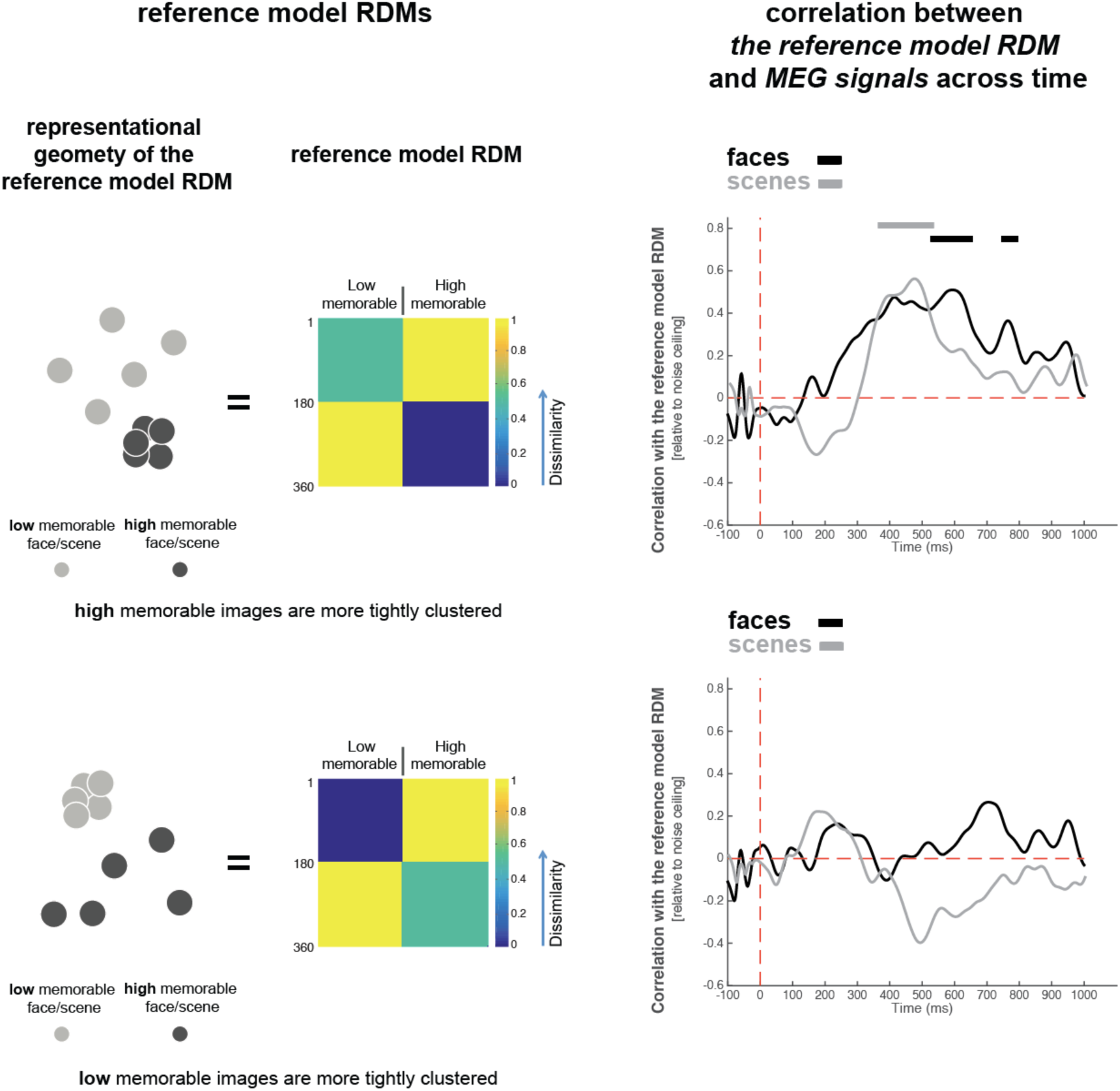
(related to Figure 2). Other candidate model RDMs for explaining the representational geometry of memorability across time. The left column (reference model RDMs) describes two different representational geometries for memorability with which the MEG RDMs are correlated across time. Patterns of correlations across time are shown on the right. The top row shows a representational space in which high memorable images are more tightly clustered than the low memorable images (the bottom row is the opposite). The RDM correlations between MEG signals and reference model RDMs are obtained similar to those in Figure 2. These results along side the result from Figure 2 (the binary memorability model RDM) shows how the representational geometry of high and low memorable images varies over time. All three model RDMs assume linear separability between high and low memorable images. Thus a significant correlation with each of them at any time-point implies that high and low memorable images could be read-out at that time with a linear classifier. Overall, the binary model RDM in Figure2 (which assumes equal tightness for clusters of high and low memorable images) and the high-memorability model RDM (tighter cluster for high memorable images–top row in this figure) both explain a significant variance of the MEG signals at different and also overlapping time-points. On the other hand, the low-memorable model RDM (tighter cluster for low memorable images–bottom row in this figure) does not explain the MEG signals at any time point, suggesting that the neural space of memorability never gets such a geometry, in which low memorable images are more tightly clustered than high memorable images. The scene memorability is equally explained well by both binary model RDM and the high-memorability model RDM starting at about 371ms after stimulus onset; later in time at t = [402ms to 622 ms] the high-memorability model RDM better explains scene memorability than the binary model RDM (two-sided signed-rank test, FDR corrected at 0.01). For faces, the binary model RDM starts to explain the MEG signals earlier (at ~ 218 ms) than the high-memorable RDM (at ~ 530 ms). These observations for both faces and scenes suggest that the high and low memorable images initially become linearly separable while each of them form clusters of almost equal tightness, and later in time the cluster of high-memorable images gets tighter than the cluster of low-memorable images.

**Figure S3.**
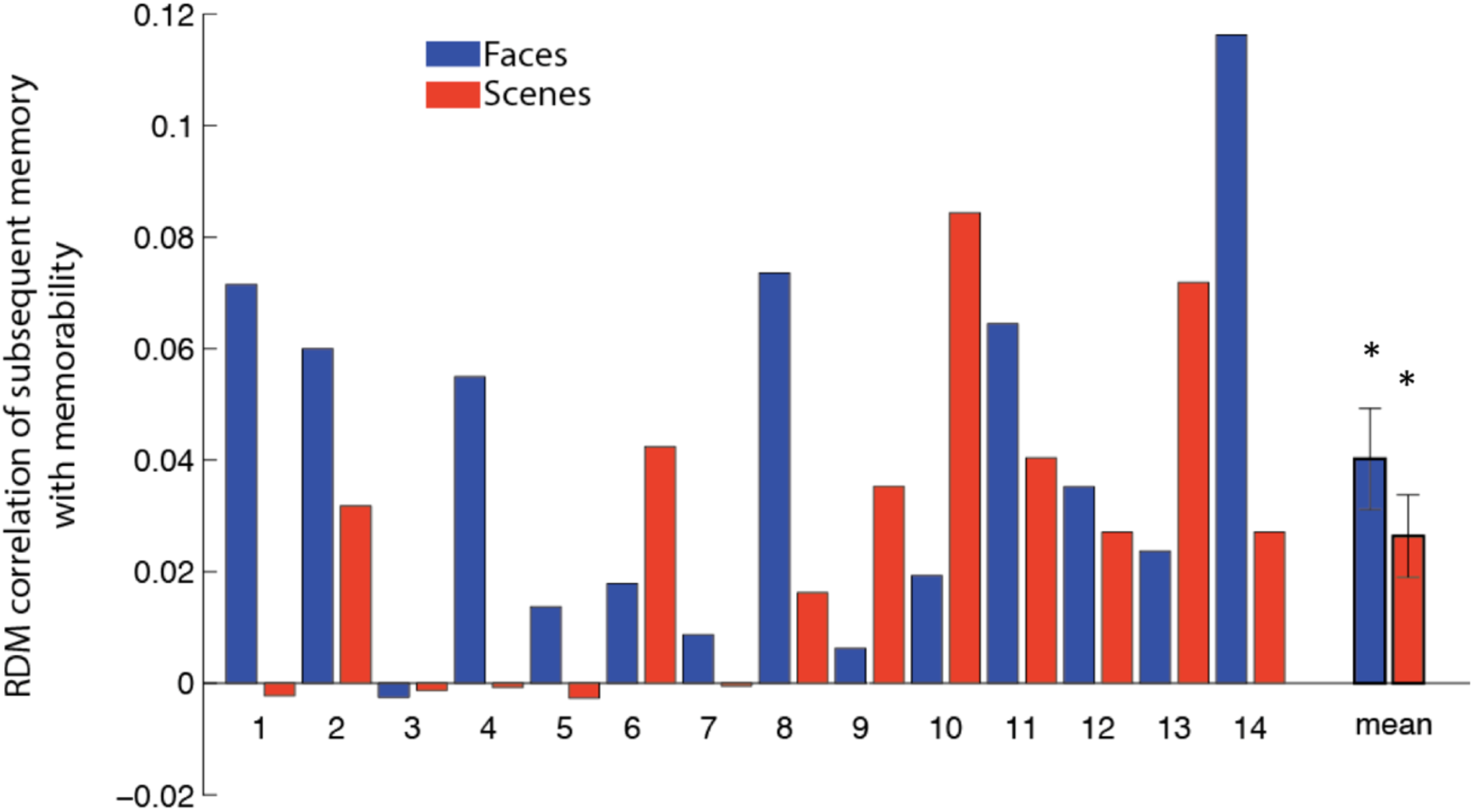
(related to Figure 2). Memorability and subsequent memory RDMs are significantly correlated. Bars indicate correlation between memorability and subsequent memory RDMs separately for each participant (N=14) and for faces and scenes. Results averaged across subjects are shown on the right. Stars above bars indicate statistical significance (N = 14, signed-rank test). Error bars are SEM.

**Figure S4.**
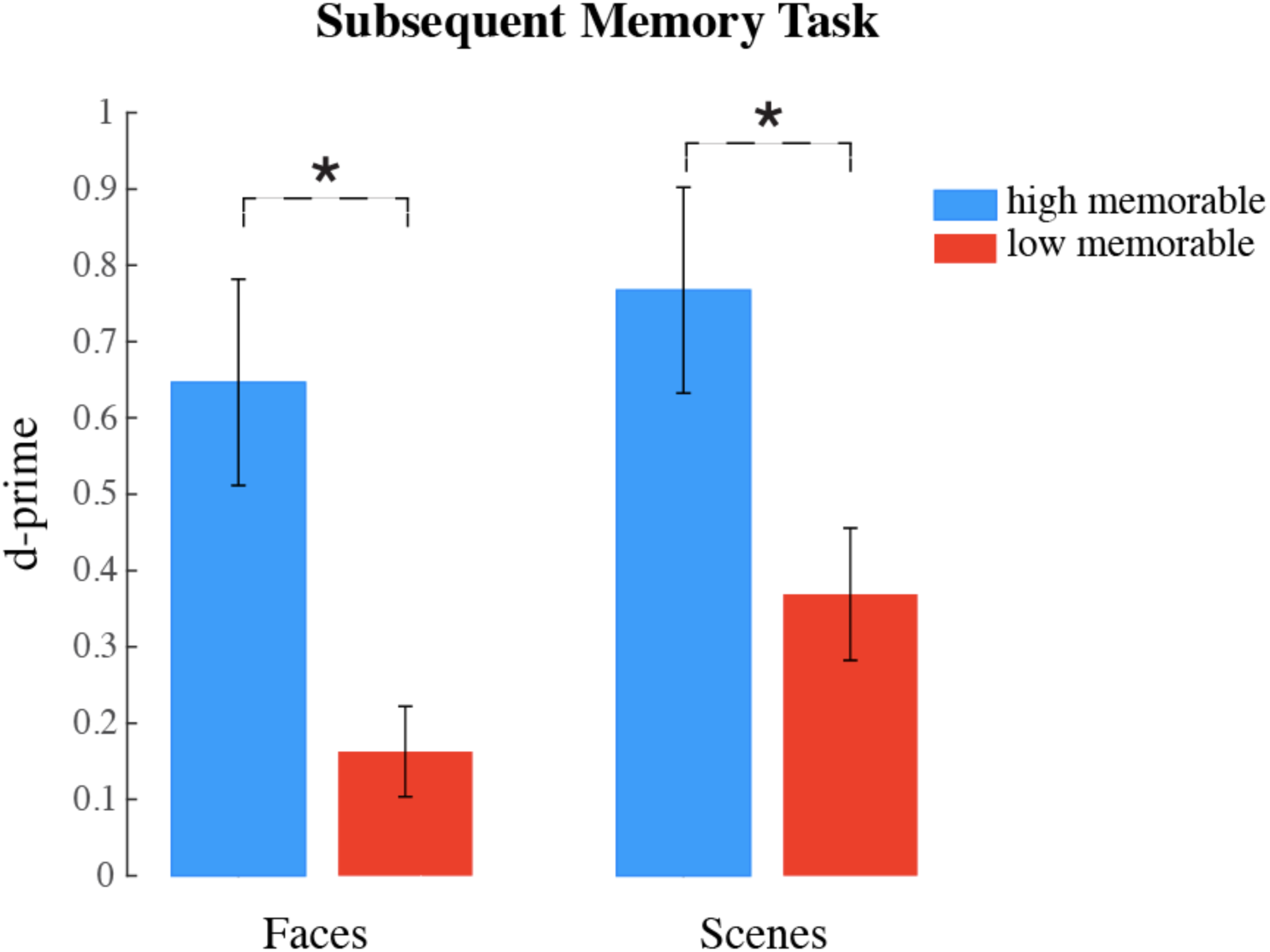
(related to ‘Experimental Procedures’). Participants significantly remembered high memorable stimuli more than low memorable stimuli in the subsequent memory test. Bars show the average performance (d’) of participants in the subsequent memory test for high (blue) and low (red) memorability images. Error bars indicate SEM; stars between bars indicate statistical significance (N=14, two-sided signed-rank test, p < 0.05). Note that the relatively low performances are expected, given that half of the images were purposely designed to be forgettable.

